# Species-specific recognition of Sulfolobales mediated by UV-inducible pili and S-layer glycosylation patterns

**DOI:** 10.1101/663898

**Authors:** Marleen van Wolferen, Asif Shajahan, Kristina Heinrich, Susanne Brenzinger, Ian M. Black, Alexander Wagner, Ariane Briegel, Parastoo Azadi, Sonja-Verena Albers

## Abstract

The UV-inducible pili system of Sulfolobales (Ups) mediates the formation of species-specific cellular aggregates. Within these aggregates, cells exchange DNA in order to repair DNA double strand breaks via homologous recombination. Substitution of the *S. acidocaldarius* pilin subunits UpsA and UpsB with their homologs from *Sulfolobus tokodaii* showed that these subunits facilitate species-specific aggregation. A region of low conservation within the UpsA homologs is primarily important for this specificity. Aggregation assays in the presence of different sugars showed the importance of *N*-glycosylation in the recognition process. In addition, the *N*-glycan decorating the S-layer of *S. tokodaii* is different from the one of *S. acidocaldarius*. Therefore, each *Sulfolobus* species seems to have developed a unique UpsA binding pocket and unique *N-*glycan composition to ensure aggregation and consequently also DNA exchange with cells from only the same species, which is essential for DNA repair by homologous recombination.

**Importance:** Type IV pili can be found on the cell surface of many archaea and bacteria where they play important roles in different processes. The Ups-pili from the crenarchaeal Sulfolobales species are essential in establishing species-specific mating partners, ensuring genome stability. With this work, we show that different *Sulfolobus* species have species-specific regions in their Ups-pilin subunits, which allow them to interact only with cells from the same species. Additionally, different *Sulfolobus* species all have unique S-layer *N*-glycosylation patterns. We propose that the unique features of each species allow the recognition of specific mating partners. This knowledge for the first time gives insights into the molecular basis of archaeal self-recognition.

## Introduction

Type IV pili (T4P) are cell surface appendages that can be found on the cell-surfaces of many bacteria and archaea (1, 2). They have been implicated in motility, secretion, DNA transformation, adhesion to surfaces and the formation of intercellular associations (3, 4). In bacteria, many examples of T4P with cellular-binding properties have been described. The major pilin subunit PilE from *Neisseria* T4P, was shown to bind endothelial cells and hemagglutinate erythrocytes whereas the *Neisseria* minor pilin PilV is essential for adherence to host cells (5–10). Additionally, major pilin PilA from *Myxococcus xanthus* binds to self-produced exopolysacharides, subsequent retraction of the T4P allows gliding motility and fruiting body formation (11, 12). T4P also form intercellular connections that are essential for conjugational exchange of DNA. For instance, PAPI-1 encoded T4P bring *Pseudomonas aeruginosa* cells in close proximity by binding to lipopolysaccharides of the recipient cells and thereby promote exchange of PAPI-1 DNA (13, 14).

In archaea, several gene clusters have been found to encode T4P-like structures (4, 15–19). The best-characterized archaeal T4P-like structure is the archaellum, which is essential for swimming motility (4, 19–21). However, little is known about the role and mode of action of archaeal non-archaellum T4P in attachment to biotic or abiotic surfaces. T4P from the thermophilic crenarchaeon *Sulfolobus acidocaldarius* (Aap: <u>a</u>rchaeal <u>a</u>dhesive <u>p</u>ili) and the euryarchaea *Haloferax volcanii* and *Methanococcus maripaludis* were shown to be involved in attachment to surfaces (22–27). However, their exact mode of binding has not been studied. Next to Aap-pili, Ups-pili can be found in Sulfolobales (<u>U</u>V inducible <u>p</u>ili of <u>S</u>ulfolobales) (28–31). These T4P assemble upon treatment of the cells with UV-stress and other DNA double strand break inducing agents. They are crucial in cellular self-interactions thereby mediating the formation of species-specific cellular aggregates (32, 33). Within these aggregates, cells are able to exchange chromosomal DNA using the Ced-system (<u>C</u>renarchaeal <u>e</u>xchange of <u>D</u>NA), suggesting a community based DNA repair system via homologous recombination (32, 34). Interestingly, the Ced-system was found to function independently of the Ups-pili, even though both systems are essential for DNA transport (35).

The *ups*-operon encodes two pilin subunits with a class III signal peptide: UpsA and UpsB (29). Deletion mutants of either *upsA* or *upsB* still form pili (though less and smaller), but do not aggregate after UV induction. The pilins are therefore both suggested to be major subunits forming mixed Ups-pili (31, 32). While the importance of Ups-pili in cellular recognition is known, the underlying molecular mechanism of the species-specific cellular aggregation of *Sulfolobus* species has not been determined.

In this study, we investigated the role of Ups-pili in species-specific aggregation on a molecular level. To this end, *in vivo* chimera mutants were constructed in which we exchanged (parts of) the genes encoding the pilin subunits UpsA and UpsB of *S. acidocaldarius* and *S. tokodaii*. By using these strains in aggregation assays and fluorescence in situ hybridization (FISH) experiments, we were able to assign a specific region of UpsA to be required for species-specific cell aggregation of archaeal cells. Furthermore, aggregation assays in the presence of different sugars suggested a role of *N*-glycosylation in cellular recognition. Glycan analysis on the thus far unstudied *S. tokodaii* S-layer showed a different *N*-glycan composition compared to that of other *Sulfolobus* species. Based on these experiments, we propose that a specific region of UpsA forms a binding site to bind species-specific *N*-glycan chains of S-layer components, thereby allowing species-specific cell aggregation and subsequent DNA exchange.

## Results

### The role of pilin subunits in species-specificity

To study the role of the Ups-pilin subunits (UpsA and UpsB) in species-specific recognition of *Sulfolobus* cells, a *S. acidocaldarius* strain was constructed in which the genomic region from the start codon of *upsA* until the stop codon of *upsB* was exchanged with the orthologous region from *S. tokodaii* (resulting in strain MW135; Figure 1A, Table 1). Upon UV-induction S. *acidocaldarius* MW135 was still found to produce Ups-pili (Figure S3), however, interestingly, it showed little to no cellular aggregation (Figure 1B). In order to test if this *S. acidocaldarius* Ups-hybrid strain was able to recognize and therefore aggregate with *S. tokodaii* cells, fluorescence in situ hybridisation (FISH) with species-specific probes was performed on mixed *S. acidocaldarius/ S. tokodaii* strains after UV-induction. A positive control with a mixture of background strain *S. acidocaldarius* MW501 (Δ*flaI*/Δ*aapF*, a strain that does not produce archaella or Aap-pili; Table 1) and *S. tokodaii*, confirmed previously observed species-specific aggregation (Figure 1C, first panel). The negative control in which a *S. acidocaldarius* Δ*upsAB* strain was mixed with *S. tokodaii*, revealed, as expected, no aggregation of the *S. acidocaldarius* Δ*upsAB* strain and normal aggregation of *S. tokodaii* (Figure 1C, second panel). Interestingly, cells from *S. acidocaldarius* MW135 interacted with *S. tokodaii* cells and thereby formed mixed species aggregates (Figure 1C, third panel).

**Table 1:** Strains used during this study.

| Strain | Background strain | Genotype | Source/ reference |
| --- | --- | --- | --- |
| *S. tokodaii* 7 |  |  | Suzuki et al. 2002 |
|  | *S. acidocaldarius* |  |  |
| MW001 | DSM639 | ΔpyrEF (91–412 bp) | Wagner et al. 2012 |
|  | *S. acidocaldarius* | *Saci upsAB::ST upsAB*, |  |
| MW135 | MW501 | ΔflaI (Δbp 1-672), ΔaapF | This study |
|  | *S. acidocaldarius* | *Saci upsA* (aa 84-98)::*ST* |  |
| MW137 | MW501 | *upsA* (aa 80-101), ΔflaI | |
| | | (Δbp 1-672), ΔaapF | This study |
| | *S. acidocaldarius* | ΔupsAB, ΔflaI (Δbp 1- | |
| MW143 | MW501 | 672), ΔaapF | This study |
|  | *S. acidocaldarius* |  |  |
| MW501 | MW001 | ΔflaI (Δbp 1-672), ΔaapF | |
|  |  |  | (53) |
|  | *S. acidocaldarius* |  |  |
| ΔAgl3 | MW001 | Δagl3 (Δsaci0423) | Meyer et al. 2011 |

**Figure 1:**
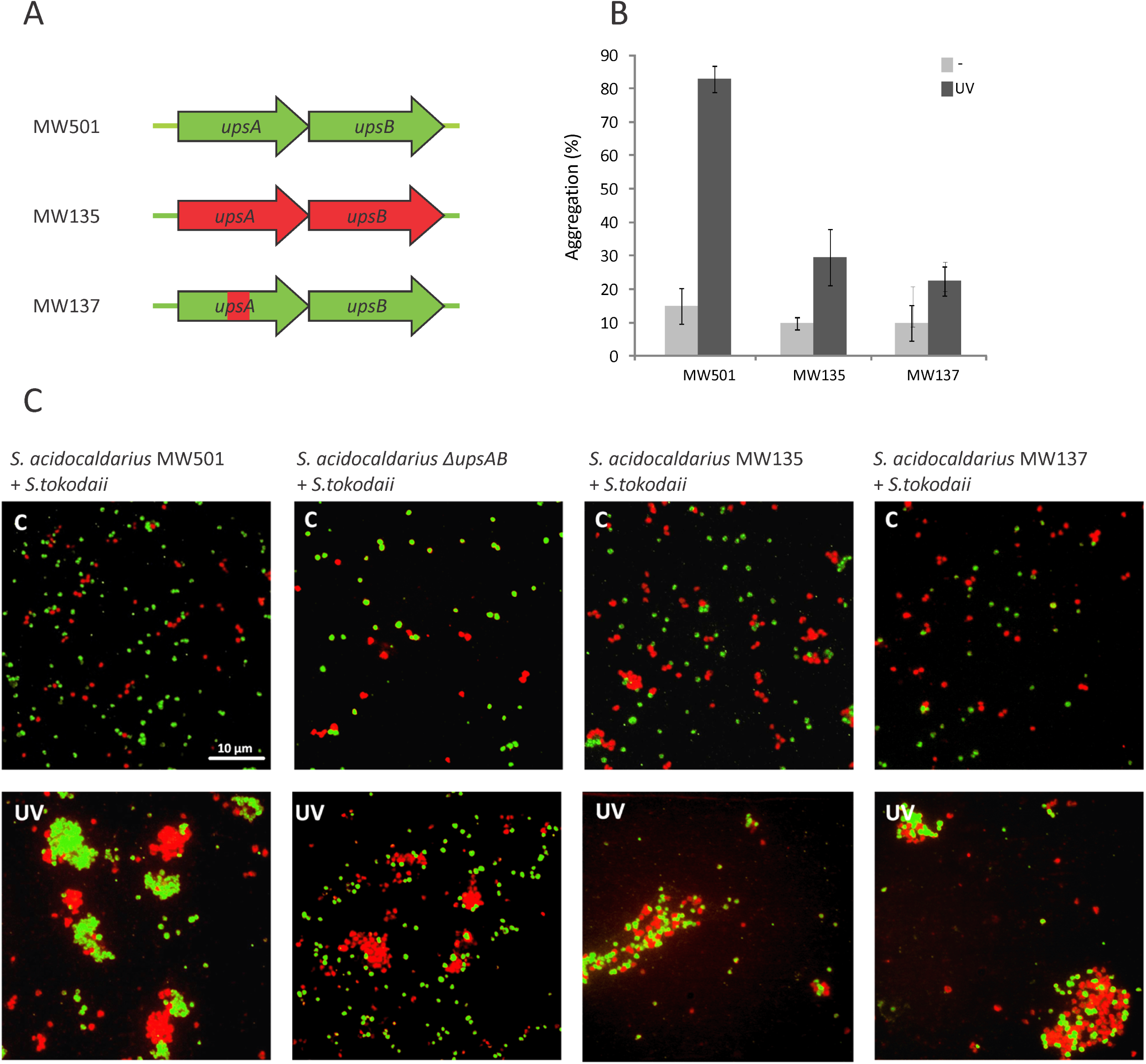
*S. acidocaldarius upsAB* mutants and their aggregation behavior. (A) (Parts of) *upsA* and *B* from *S. acidocaldarius* or MW501 (Δ*flaI*/ Δ*aapF*) (green) were replaced with the same regions from *S. tokodaii* (red), resulting in MW135 (exchange from start codon of *upsA* until stop codon of *upsB*) and MW137 (*Saci upsA* aa 84-98::*ST upsA* aa 80-101) (see also Figure S1). (B) Quantitative analysis of UV-induced cellular aggregation of mutants shown in A. Percentage of cells in aggregates 3h after induction with or without 75 J/m^2^ UV (dark or light grey, respectively). (C) Aggregation behavior of mixtures of *S. tokodaii* (red) with different *S. acidocaldarius* mutants (green) after treatment with UV-light (UV). Untreated cells were used as a control. Mutants used for this experiment were: MW501 (wt *upsAB*), MW143 (Δ*upsAB*), MW135 and MW137. FISH labeled cells were visualized with fluorescence microscopy. *Scale bar* 10 µm.

To find putative species-specific regions in the pilin subunits involved in species-specific recognition, alignments were made using UpsA and UpsB amino acid sequences, from several Sulfolobales (Figure S1). Additionally, the relationship between UpsA and UpsB homologs was studied by creating a phylogenetic tree (Figure S2, Supplementary results and methods). A region with low conservation was revealed in UpsA (Figure S1, amino acid 84-98 for *S. acidocaldarius*, red box). To test whether this region plays a role in cell-cell recognition, a strain was constructed in which only the region of low conservation in *S. acidocaldarius* UpsA (amino acid 84-98) was exchanged with the corresponding part from *S. tokodaii* UpsA (amino acid 80-101) (MW137 Figure 1A, Table 1). Similar to what was observed for the *S. acidocaldarius* mutant in which *upsA* and *upsB* were exchanged completely (MW135), *S. acidocaldarius* MW137 showed pili formation (Figure S3) but showed little to no aggregation with itself (Figure 1B). Instead, it was found to aggregate with *S. tokodaii* (Figure 1C, fourth panel). This observation strongly suggests that the non-conserved region (exchanged in MW137) defines the species-specificity during cellular aggregation.

### The role of glycosylation in species-specificity

The fact that *Sulfolobus* Ups wild-type strains are able to form mating pairs with Ups-deletion strains (32), suggests that factors, other than Ups-pili, play a role in species-specific recognition. The surface proteins of Sulfolobales are heavily glycosylated (36, 37). We therefore suggested that Ups-pili might recognize glycosylated proteins and thereby initiate cellular interactions. To confirm this hypothesis, UV induced aggregation assays were performed in the presence of monosaccharides that are also part of the *S. acidocaldarius N*-glycan chain (Glc_1_Man_2_GlcNAc_2_QuiS, containing glucose, mannose, *N*-acetylglucosamine and the *Sulfolobus*-specific sulfoquinovose residues) (37) (Figure 2). The addition of *N*-acetylgucosamine or glucose did not result in altered cellular aggregation (Figure 2A and B); however, in the presence of mannose, cell aggregates were significantly smaller (Figure 2B). This suggests that the mannose molecules partially saturate the binding sites of the Ups-pili and thereby inhibit interactions between pili and the glycan chains on the S-layer of the host cell resulting in reduced aggregation.

**Figure 2:**
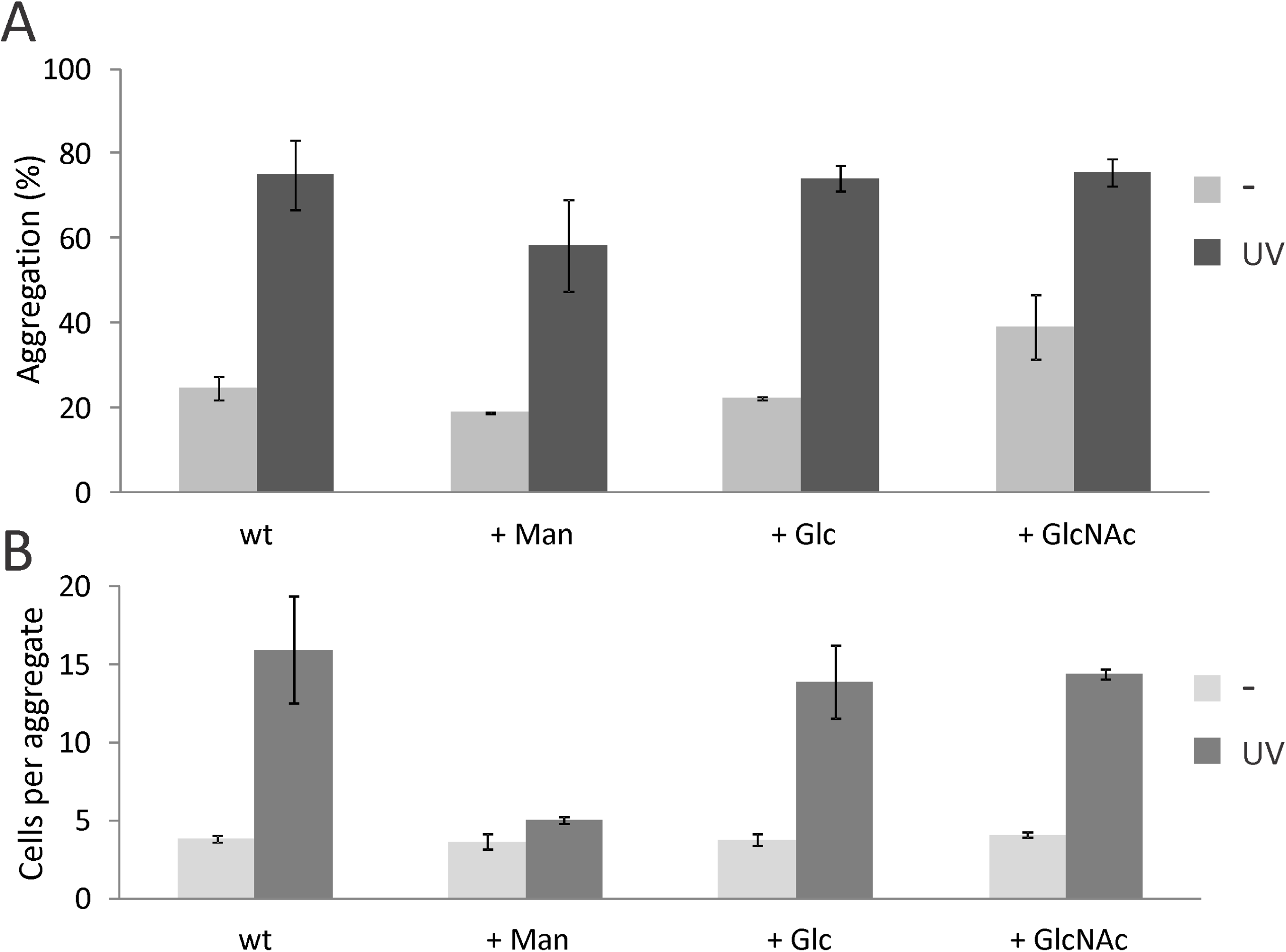
UV-induced aggregation of *S. acidocaldarius* MW001 upon addition of 20 mM mannose, glucose or *N*-acetylglucosamine. (A) Percentage of cells in aggregates. (B) Average sizes of formed aggregates. Light grey bars represent non-induced cells and dark grey bars represent cells induced with 75 J/m^2^ UV.

### Defining the glycosylation pattern of *S. tokodaii* S-layer proteins

Our hypothesis that S-layer glycosylation is important for species-specificity suggests that different *Sulfolobus* species have different glycosylation patterns. So far, the glycan structure of *S. tokodaii i*s unknown. To analyze the glycan structures on the S-layer of *S. tokodaii, N*-glycans were released from isolated S-layer by hydrazinolysis. Using MALDI/TOF-MS profiling, one main *N*-glycan species and two other low abundant species could be identified in both positive (Figure S4A) and negative ion mode (Figure S4B). The structures of *N*-glycans were proposed based on mass-to-charge ratio of each *N*-glycan ions observed (Figure S4, Table 2) as well as its MS/MS fragmentation pattern (Figure 3). The three *N*-glycan species could be identified as QuiS_1_Hex_4_HexNAc_2_, QuiS_1_Hex_3_HexNAc_2_, and QuiS_1_Hex_4_HexNAc_1,_ respectively (Table 2). To determine the linkages between the sugars in the deduced *N*-glycan species, linkage analysis (38) was performed on the permethylated *N*-glycans released from S-layer proteins. The various types of linkages observed on each monosaccharide and their relative abundances on the *N*-glycans are shown in the Figure S5. The most plausible position of this linkage in the glycan chain can be observed on the right-side column in Figure S5. Based on this linkage information, MS^n^ determination of glycan branching (Figure 3), and the glycan masses (Figure S4), the *N*-glycan glycoforms and their isomers were deduced (Figure S6). Figure 4 schematically shows the most prominent glycan structures from *S. acidocaldarius* (37), *S. solfataricus* (39) and *S. tokodaii* (this study). In agreement with our hypothesis, the core of these structures are similar, whereas the outer saccharides differ. A typical sulfated glycan is present in all three *Sulfolobus* glycan structures. Using LC-MS profiling on the tryptic digest of SlaA and SlaB, several different glycopeptides could indeed be observed (Supplementary results and methods, Figure S7 and Figure S8 respectively).

**Table 2.** List of *N*-linked glycans released from S-layer glycoprotein from *S. tokodaii* detected by MALDI/TOF-MS. Abbriviations: QuiS: sulfoquinovose, Hex: hexose, HexNAc *N*-acetyl hexosamine.

| Permethylated mass (m/z)<sup>1</sup> | Text description of structures | Percentage of glycans |
| --- | --- | --- |
| 1406 | QuiS<sub>1</sub>Hex<sub>4</sub>HexNAc<sub>1</sub> | 3.92 |
| 1447 | QuiS<sub>1</sub>Hex<sub>3</sub>HexNAc<sub>2</sub> | 7.09 |
| 1651 | QuiS<sub>1</sub>Hex<sub>4</sub>HexNAc<sub>2</sub> | 88.99 |

<sup>1</sup>All masses (mass+ 2Na - H) are single-charged.
<sup>2</sup>Calculated from the area units of detected *N*-linked glycans.

**Negative ion mode**
| Permethylated mass (m/z)<sup>1</sup> | Text description of structures | Percentage of glycans |
| --- | --- | --- |
| 1360 | QuiS<sub>1</sub>Hex<sub>4</sub>HexNAc<sub>1</sub> | 1.27 |
| 1401 | QuiS<sub>1</sub>Hex<sub>3</sub>HexNAc<sub>2</sub> | 1.82 |
| 1605 | QuiS<sub>1</sub>Hex<sub>4</sub>HexNAc<sub>2</sub> | 96.91 |

<sup>1</sup>All masses (mass - H) are single-charged.
<sup>2</sup>Calculated from the area units of detected *N*-linked glycans.

**Figure 3:**
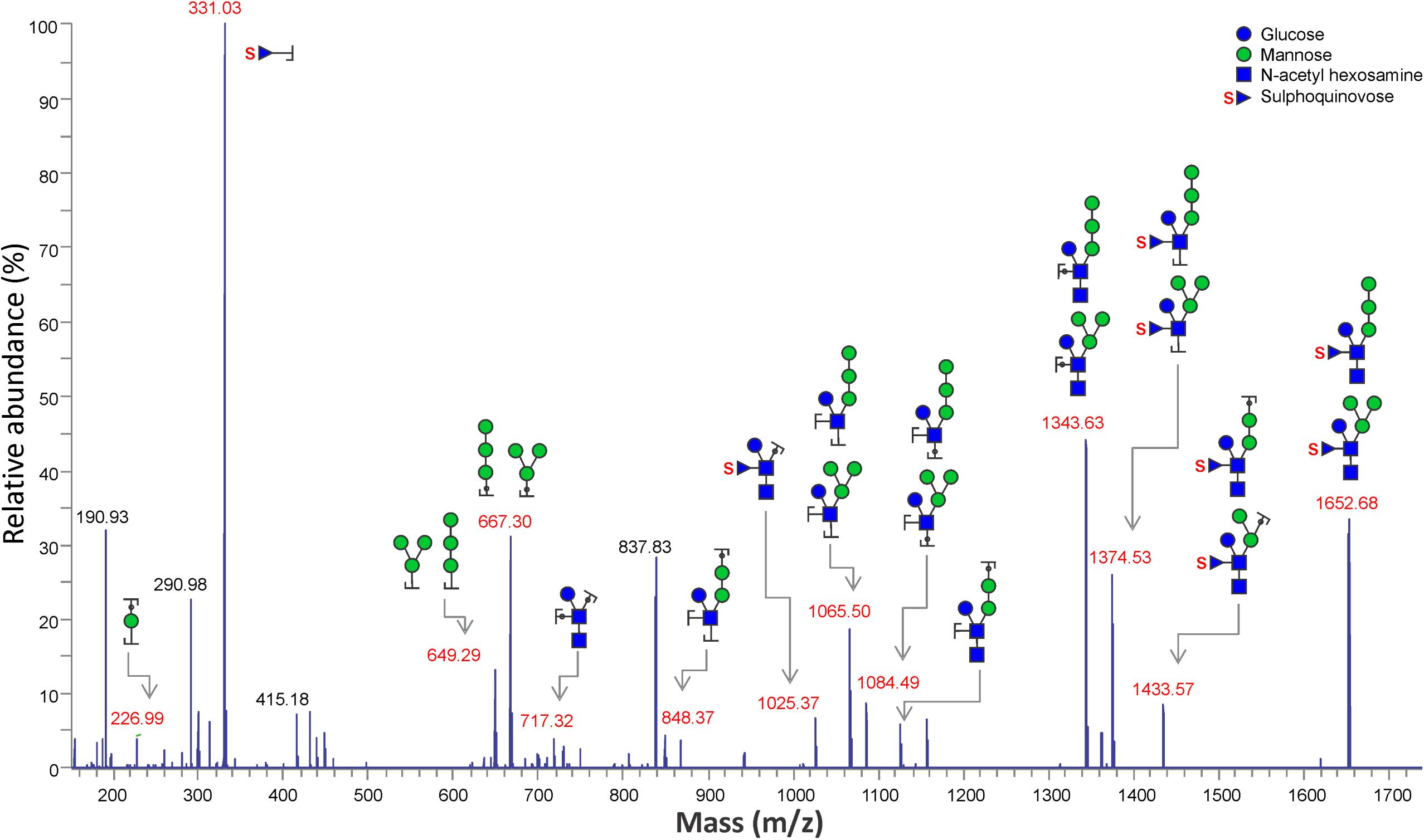
HCD MS^2^ spectra of heptasaccharide (m/z - 1651.7, Figure S4a) released from the S-layer proteins from *S. tokodaii* by hydrazinolysis.

**Figure 4:**
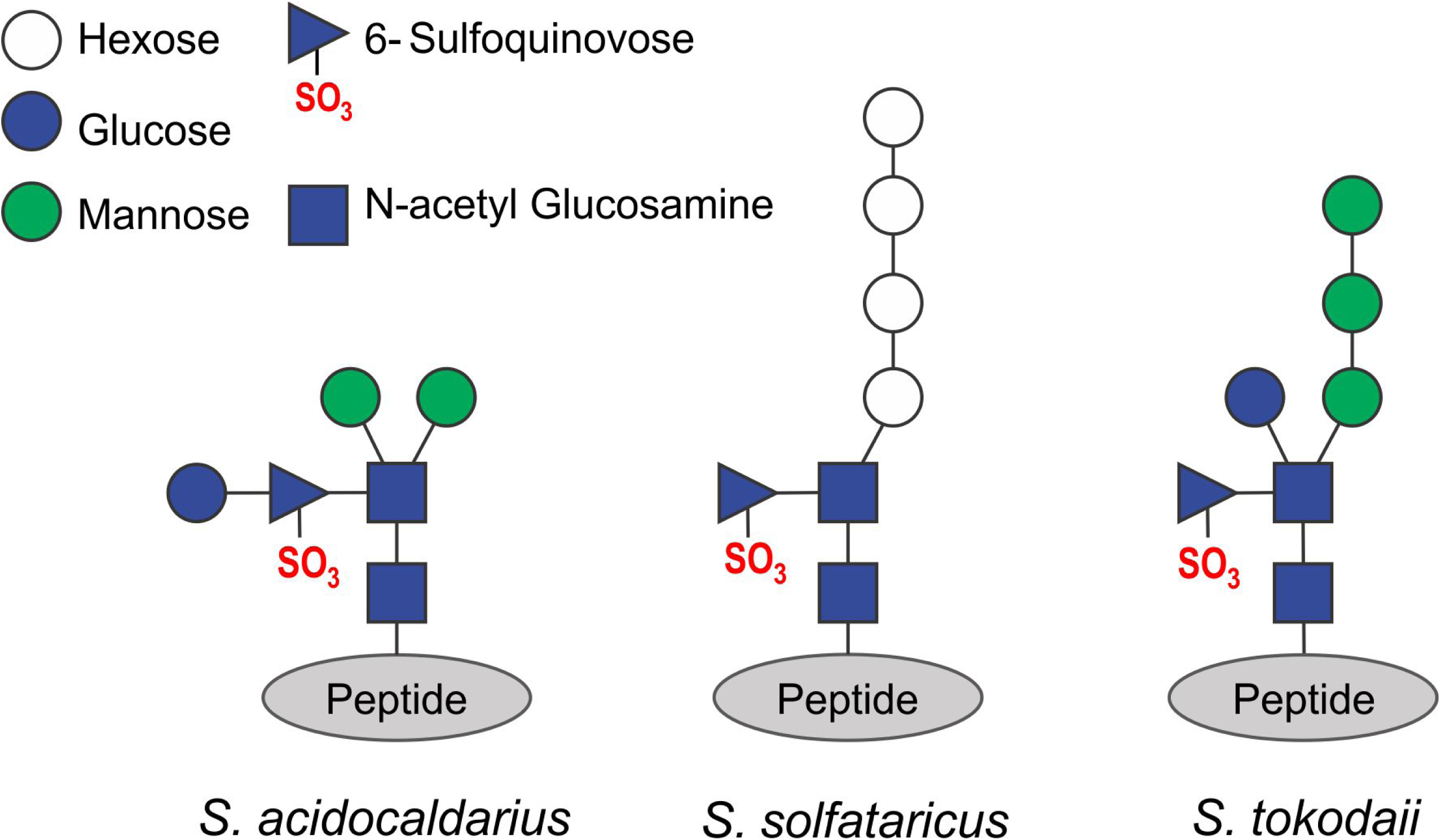
Structure of the glycan trees present on the S-layer of *S. tokodaii* in comparison to those from *S. acidocaldarius* (37) and *S. solfataricus* (39).

### Determination of the binding site in UpsA

We know that a *S. acidocaldarius* mutant in which both Ups-pilin subunits are deleted, does not aggregate upon UV-stress (31). Here we could successfully complement this phenotype by expressing the *upsAB* genes from a maltose inducible plasmid (Figure 5, Δ*upsAB* + *upsAB*). Using site directed-mutagenesis on this plasmid, we moreover created point mutations within the above-described region of interest of UpsA (black squares in Figure S1): D85A, N87A, N94A and Y96A. All mutants still produced pili upon UV induction (Figure S8). Interestingly, when expressing UpsA in which the poorly conserved residues D85 or Y96 were mutated to alanine, respecitvely, UV induced aggregation was significantly reduced. On the other hand, mutation of the conserved N87 or N94 showed wild-type aggregation (Figure 5). These results suggest that the region of low conservation within UpsA is specifically adapted to the glycan structure of the same species in order to ensure species-specific aggregation only.

**Figure 5:**
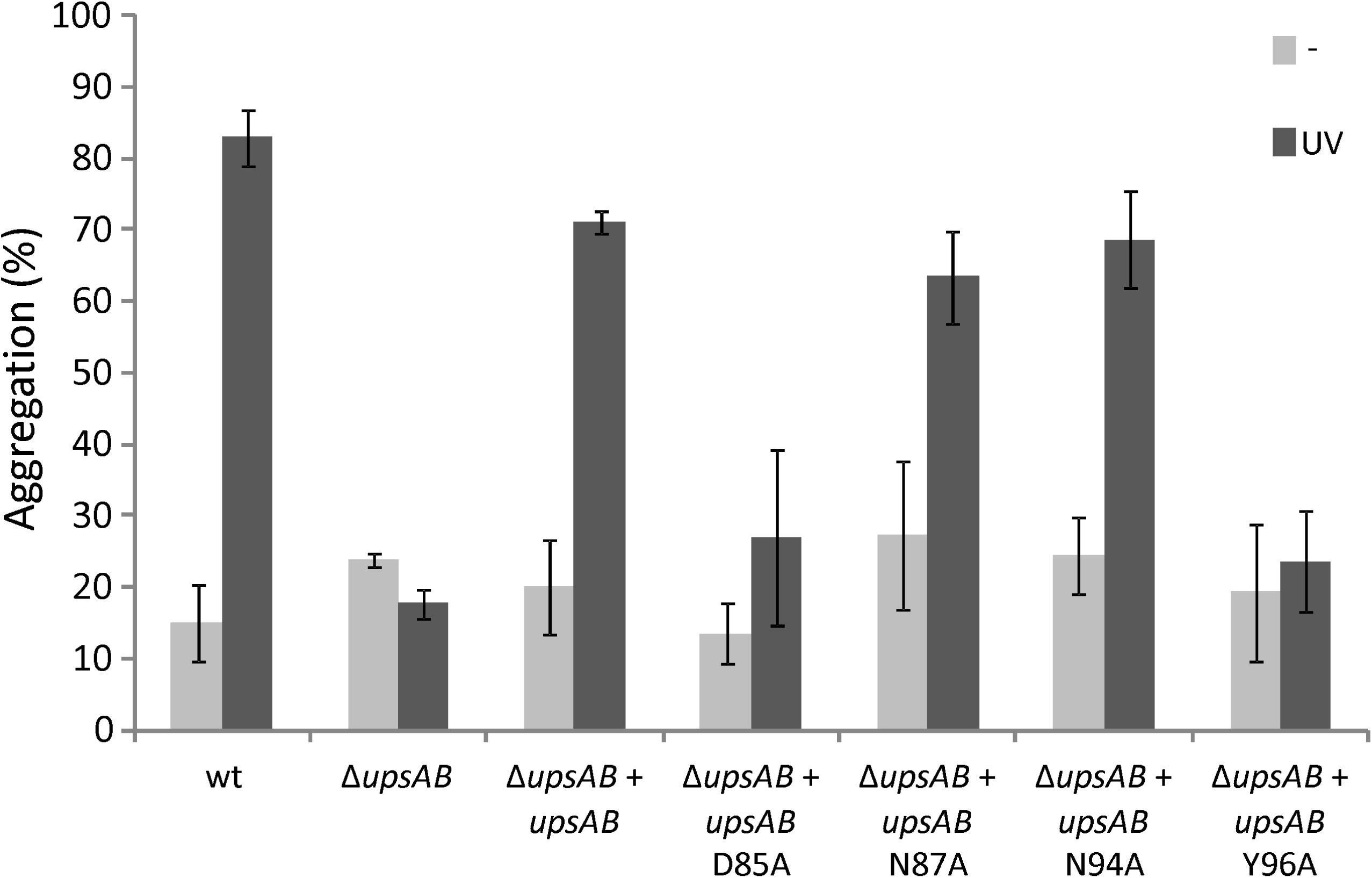
UV induced cellular aggregation of *S. acidocaldarius* Δ*upsAB* complementation strains. A *S. acidocaldarius* Δ*upsAB* mutant (MW143) was complemented with maltose inducible plasmids carrying *upsAB* or *upsAB* with a D85A, N87, N94A or Y96A mutation in UpsA (see also Figure S1). Percentage of cells in aggregates 3h after induction with or without 75 J/m^2^ UV (dark or light grey, respectively).

## Discussion

Both bacterial and archaeal T4P have shown to be essential for surface-adherence. Given the fact that bacterial T4P are strongly related to pathogenicity, their mode of binding has primarily been studied for pathogenic bacteria such as *P. aeruginosa, Vibrio cholerae, Myxococcus, Neisseria* and Enteropathogenic *E. coli* species. However, also non-pathogenic bacteria and archaea encode several T4P involved in adhesion, which are studied in far less detail. The Crenarchaeal Sulfolobales encode three types of T4P: archaella, involved in swimming motility (20); Aap-pili, involved in attachment to diverse surfaces (24, 25); and Ups-pili, mediating intraspecies cellular aggregation and DNA exchange (29, 31, 32, 35). During this study, we have examined the role that Ups-pili play in the formation of *Sulfolobus* mating partners. In particular, we focused on the role that pilin subunit UpsA plays in cell-recognition.

The Ups-pilus is formed by two pilin subunits UpsA and UpsB, which are both thought to be major pilin subunits that build up mixed pili structures (31). We revealed that UpsA is involved in species-specific cellular interactions and we were able to alter this specificity by exchanging (parts of) the pilin subunit with that of another species (Figure 1). The binding of bacterial T4P to other cells is often based on pilin-sugar interactions. Surface exposed glycans can be found on cells from all domains of life where they display an enormous range of different structures that are often highly specific to certain species (40). Glycans are therefore perfect anchors to bind specific host- or partner cells. In *Saccharomyces cerevisiae*, surface exposed lectins can bind to surface exposed sugars in a calcium-dependent manner, thereby forming cellular aggregates, a process which is called flocculation (41). This behavior can be inhibited by saturating the binding of the lectins through addition of loose sugars to the medium (42) (Figure 2). In similar experiments with S. *acidocaldarius* we found that mannose has an inhibiting effect on UV-induced cellular aggregation. Since two outer mannose residues are present in the *S. acidocaldarius N*-glycan tree, binding of Ups-pili to this side of the glycans tree is probable. When analyzing the *N*-glycans of *S. tokodaii*, we could indeed find differences in this part of the *N*-glycan structure when compared to that of *S. acidocaldarius* (37) and S. *solfataricus*. (39) (Figure 4). As observed for Eukarya (43), the core or the glycan structure is similar in all three species, whereas the outer residues differ. Our results thereby suggest that UpsA contains a specific binding pocket that is able to bind specific sugar moieties of the *N*-glycans presented on the S-layer of distinct *Sulfolobus* species (Figure 6).

**Figure 6:**
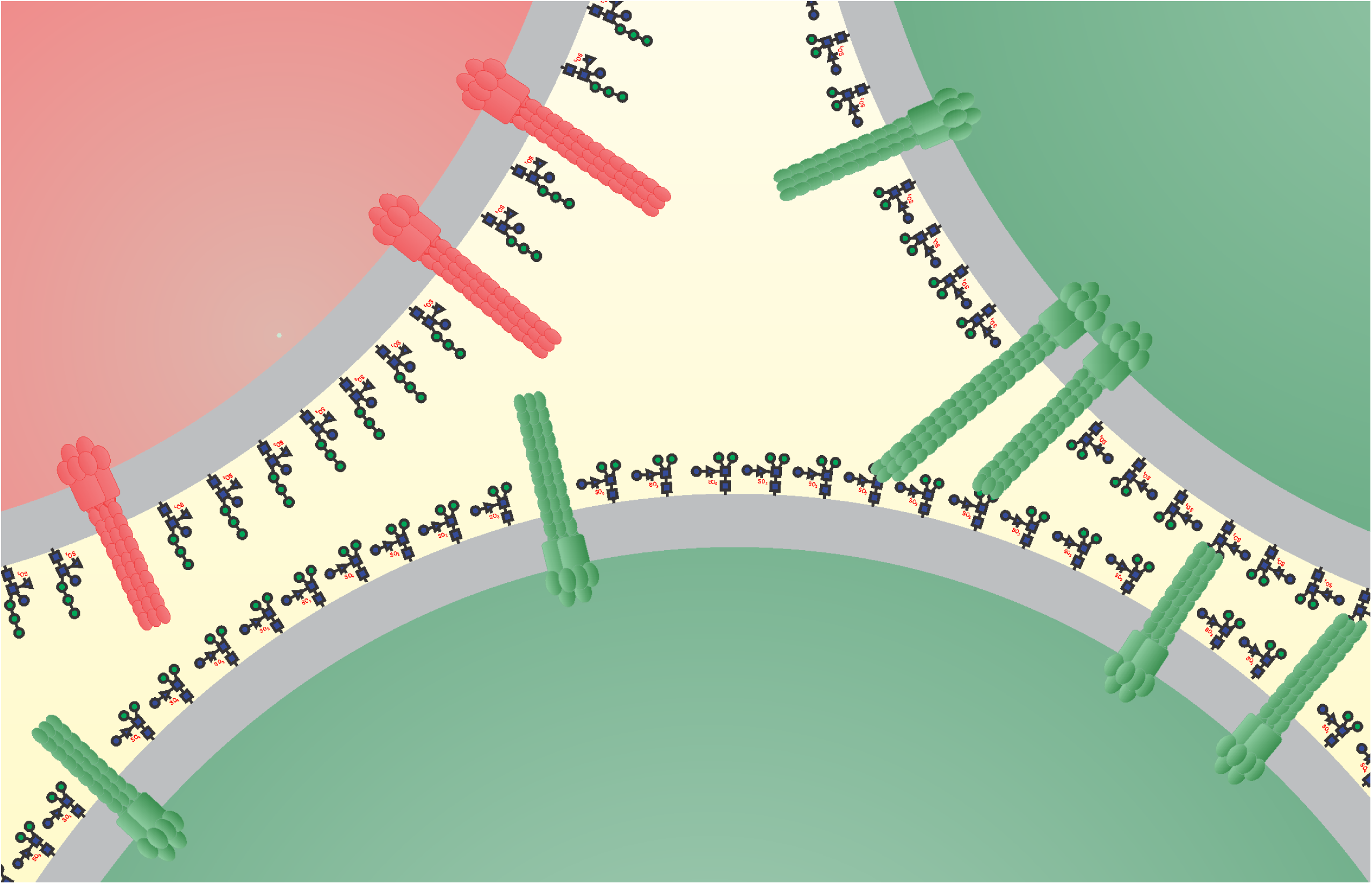
Proposed model species-specific interactions between Ups-pili and *N*-glycosylated S-layer of Sulfolobales. Ups-pili of *S. acidocaldarius* (green) only form interactions with the *N*-glycan of the same species and not with that of other species *S. tokodaii* (red).

Among the euryarchaeal *Haloferax* species, glycosylation was found to be essential for cell fusion (44), emphasizing the importance of glycosylation in Archaeal cellular recognition in general. It is unclear if pili or other types of lectin molecules are involved in cellular interactions that initiate *Haloferax* fusion events. Similar to our findings, different *Haloferax* species are also known to be differentially glycosylated (45) leading to semi-specific cell-cell recognition (44). Cell fusion between different *Haloferax* species could also be observed but with far lower efficiency (46). In addition, under different environmental conditions, *Haloferax* glycosylation patterns change, leading to more or less favorable *N*-glycans for mating (47). One could envision that low frequency interactions between different *Sulfolobus* species also occur and might occasionally lead to horizontal gene transfer (34), thereby affecting speciation. In a single hot spring in Kamchatka (Russia) two different groups of *Sulfolobus islandicus* strains were found to be present. Despite their coexistence, it was postulated that *S. islandicus* species mainly exchange DNA within these groups (48). It is likely that *N*-glycan patterns and Ups-pili between the species are different serving as a barrier to gene transfer. This behavior might be seen as the two groups diverging into different species.

The Ced-system that is involved in DNA transfer among Sulfolobales can also be found in several crenarchaea that do not encode Ups-pili (34, 35), it is therefore likely these species have developed a different mechanism to initiate cellular interactions. Given the importance of glycosylation in cell-cell interactions in both euryarchaeal *Haloferax* and crenarchaeal *Sulfolobus* species, glycosylation most likely also plays a role in these interactions.

This study has given molecular insights in the cellular recognition mechanism of the previously described crenarchaeal Ups-system (29, 31, 32). Our current model suggests that upon DNA damage, Ups-pili are formed; the UpsA pilin subunits contain a species-specific glycan binding pocket in pilin subunit UpsA that can only bind glycans presented on cells from the same species (Figure 6). This system allows the formation of species-specific cellular connections prior to DNA-exchange via the Ced-system (35). In that way, only DNA from the same species is exchanged and used for DNA repair via efficient homologous recombination. This proposed cellular recognition mechanism in Sulfolobales restricts exchange of genomic DNA to cells from the same species, thereby playing an important in role genome integrity and the maintenance of species. Co-evolution of *N*-glycosylation and pilin subunit UpsA might play an important role in speciation.

## Experimental procedures

### Culture conditions

*Sulfolobus acidocaldarius* strains and derived mutants (Table 2) were grown aerobically at 75 °C in basic Brock medium (49), supplemented with 0.1% NZ amine, 0.2% dextrin and 20 µg/ml uracil and adjusted to pH 3.5 with sulfuric acid. For solid media the medium was supplemented with 1.5% gelrite. Plates were incubated for 5-6 days at 76°C. *E. coli* competent cells DH5α and ER1821 (NEB) used for respectively cloning and methylation of plasmid DNA were grown in LB medium (10 g/l tryptone; 5 g/l yeast extract; 10 g/l NaCl) at 37°C supplemented with the appropriate antibiotics. Growth of cells was monitored by optical density measurements at 600 nm.

### Deleting, exchanging and complementation of genes in *S. acidocaldarius*

To construct deletion and pilin exchange mutants; up- and downstream flanking regions of the genes of interest (approximately 600 bp) were amplified with primers listed in Table S1. Overlap PCR was performed to connect the up- and downstream fragments. To replace (parts of) *upsA* and *upsB* from *S. acidocaldarius* with their homologues from *Sulfolobus tokodaii*, synthetic DNA was ordered (GenScript) consisting out of *S. acidocaldarius upsAB* flanking regions and (parts of) *S. tokodaii upsAB* genes (Table S1). The PCR product and synthetic DNA fragments were subsequently cloned into pSVA406, resulting in the plasmids listed in Table 1. The plasmids were methylated in *E. coli* ER1821 containing pM.EsaBC4I (NEB) (50) and transformed into *S. acidocaldarius* MW501 (Δ*fla/*Δ*aap*) (Table 1) (51). This background strain lacks Aap-pili and archaella, allowing easy EM analysis. Integrants were selected on plates lacking uracil and grown in 24-well plates for 2 days in the same medium. Subsequently cultures were plated and grown for 5 days on second selection plates containing uracil and 100 µg/ml 5-FOA to select for clones in which the plasmid looped out by homologous recombination. Obtained colonies were tested by PCR for successful deletion/replacement of the genes. Correctness of strains was confirmed by DNA sequencing. Strains that were made during this study are listed in Table 1.

For complementation of a Δ*upsAB* mutant (MW143), the DNA region comprising *upsA* and *B* was amplified using primers listed in Table S1 and cloned into pSVA1450 under control of a maltose inducible promoter resulting in plasmid pSVA1855 (Table S1). This plasmid was subsequently used to as a template to introduce point mutations into *upsA* (D85A, N87A, N94A and Y96A) (Table 1) using two overlapping primers per mutation (Table S1). Resulting plasmids were then transformed via electroporation into MW143 as described previously (51). Cultures were grown without the addition of uracil. Expression of (mutated) UpsA and B was induced by addition of 0.2% maltose.

### UV treatment, aggregation assays

UV light treatment was performed as described in (29); 10 ml culture (OD_600_ 0.2-0.3) was treated with a UV dose of 75 J/m^2^ (254 nm, Spectroline, UV crosslinker) in a plastic petri dish. For FISH experiments *S. acidocaldarius* and *S. tokodaii* were first mixed in equal amounts. For complementation the Δ*upsAB* strain, expression of UpsAB (derivatives) was additionally induced with 0.2% maltose. Afterwards cultures were put back at 76°C for 3 h. Samples taken at different time points were analyzed with phase contrast microscopy. To quantify aggregated cells after induction with UV, 5 µl of cell culture (diluted to OD 0.2) was spotted on a microscope slide covered with a thin layer of 2% agarose in Brock minimal medium. Cells were visualized with phase contrast microscopy (Zeiss, Axio Observer.Z1). Free and aggregated cells (≥ 3) were counted for at least three fields per strain using ImageJ cell counter. Percentages of cells in aggregates were subsequently calculated.

### Fluorescence *in situ* hybridization (FISH)

For FISH experiments, 10 µl of a mixed UV induced (described above) culture was spotted and dried on a glass slide. To fix the cells, 10 µl of 37% formaldehyde was spotted on top of the cells and incubated for 20 min at room temperature. Afterwards formaldehyde was removed and the cells were washed for 10 min with a drop of 1x PBS. Glass slides were subsequently dried at room temperature. Cells were permeabilized by incubating the slides 3 min in 50, 80 and 96% ethanol, respectively. After drying the slides, 10 µl of hybridization buffer (900 mM NaCl, 20mM Tris HCl pH 8.0, 10% formamid) mixed with 50 ng/µl FISH probes (for *S. acidocaldarius* and *S. tokodaii*, Table S1) was spotted on the cells. Slides were incubated in the dark at 46 °C for 1.5 h for hybridization. Subsequently the cells were washed by incubating the slides for 10 min in wash buffer (450 mM NaCl, 20 mM Tris HCl pH 8.0) at 48°C. Slides were then dipped in ice cold water and dried. For microscopy, 1x PBS was spotted on the cells and a coverslip was added. Cells were examined using fluorescence microscopy (Zeiss, Axio Observer.Z1).

### S-layer isolation

A cell pellet from a 50 ml *S. tokodaii* str. 7 culture with an OD_600_ of about 0.6 was resuspended and incubated whilst shaking (500 rpm) for 60 min at 37°C in 30 ml of buffer (10 mM NaCl, 20 mM MgSO_4_, 0.5% sodium lauroylsarcosine, pH 5.5). Samples were centrifuged for 45 min in an Avanti J-26 XP centrifuge (Beckman Coulter) at 21,000 g (rotor JA-25.50), yielding a brownish tan pellet. The pellet was resuspended and incubated for 30 min at 37°C in 1 ml of buffer A. Subsequent centrifugation for 20 min (tabletop centrifuge at maximum speed), yielded in a translucent tan pellet containing S-layer proteins. Purified S-layer proteins were washed a few times with water and then stored in water at 4°C.

### Electron microscopy analysis

Ups-pili on *S. acidocaldarius* cells were visualized with TEM. Cells were negatively stained with 2% uranyl acetate on carbon-coated copper grids. Transmission electron microscopy images were recorded using the Talos L120C (Thermo Scientific™) microscope equipped with a 4k × 4k Ceta CMOS camera. Acceleration voltage was set to 120kV and magnification to 2.27Å/ pixel.

## Acknowledgements

AS, IMD and PA were supported in part by the National Institutes of Health grants 1S10OD018530 and P41GM10349010 to the Complex Carbohydrate Research Center. SB was supported by a postdoctoral fellowship from the German Academy of Sciences Leopoldina. We thank Małgorzata Ajon (University of Groningen) for technical support.

## Supplementary material

**Supplementary results and methods:** describing the phylogenetic analysis of UpsA and UpsB from different Sulfolobales and the *N*-glycan analysis of the glycosylated S-layer of *S. tokodaii* S-layer proteins SlaA and SlaB.

**Table S1:** Plasmids and primers used during this study.

**Figure S1:** Alignments of UpsA from different Sulfolobales. The class III cleavage site, cleaved by PibD is depicted by a red line. The red box in UpsA indicates the less conserved region. Shown are UpsA amino acid sequences from *Sulfolobus acidocaldarius* DSM 639 *(Saci), Sulfolobus tokodaii* Str. 7 *(ST), Sulfolobus solfataricus* P2 *(Sso), Stygiolobus azoricus (Staz), Metallosphaera cuprina* Ar-4 *(Mcup), Metallosphaera hakonensis* DSM 7519 (*Mhak*), *Metallosphaera sedula* DSM5348 *(Msed)*, and *Metallosphaera yellowstonensis* MK1 (*Myel*).

**Figure S2:** Maximum-likelihood phylogenetic tree of UpsA and UpsB homologs from different archaeal species (*Sulfolobus acidocaldarius* strains: DSM 639, N8, Ron12/I, SUSAZ; *Sulfolobus solfataricus* strains: P2, 98/2, P1; *Sulfolobus islandicus* strains: REY15A, HVE10/4, M.16.4; *Sulfolobus tokodaii* str. 7; *Stygiolobus azoricus; Metallosphaera sedula* DSM5348; *Metallosphaera cuprina* Ar-4; *Metallosphaera hakonensis* DSM 7519; *Metallosphaera yellowstonensis* MK1). Branch numbers represent bootstrap values above 80% (100 replicates).

**Figure S3:** Transmission electron micrographs of UV induced *S. acidocaldarius* mutants (upper panel) and expression strains (lower panel). Upper panel: Ups-pili of *S. acidocaldarius* MW501 (Δ*flaI*/ Δ*aapF*), MW135 (exchange *S. acidocaldarius upsAB* genes with those of *S. tokodaii* from start codon of *upsA* until stop codon of *upsB*) and MW137 (*Saci upsA* aa 84-98::*ST upsA* aa 80-101). Lower panel: UV induced *S. acidocaldarius* expression strains. Wildtype and mutated *upsAB* genes (black squares in Figure S1) were expressed in a Δ*upsAB/*Δ*flaI*/Δ*aapF* strain (MW143). The following maltose-inducible expression plasmids were used: pSVA1855 expressing wildtype *upsAB;* pSVA1860, expressing *upsAB* with a D85A mutation in *upsA*; pSVA1860, expressing *upsAB* with a D85A mutation in *upsA*. pSVA1861, expressing *upsAB* with a N87A mutation in *upsA;* pSVA1862, expressing *upsAB* with a N94A mutation in *upsA;* pSVA1863, expressing *upsAB* with a Y96A mutation in *upsA*. Scale bar 100 nm.

**Figure S4:** (A) MALDI MS spectra of *N*-glycans released from the S-layer protein from S. *tokodaii* by hydrazinolysis observed (positive ion mode). (B) MALDI MS spectra of *N*-glycans released from the S-layer protein from S. *tokodaii* by hydrazinolysis observed (negative ion mode). Structures are assigned based on MS/MS analysis.*\*During hydrazinolysis a fraction of glycans gets derivatized by hydrazine reagent*.

**Figure S5:** Glycosyl linkages of monosaccharides of *N*-glycans from the S-Layer proteins of *S. tokodaii* were determined by GC-MS analysis using the PMAA (Partially Methylated Alditol Acetate) method.

**Figure S6:** Different possible glycoforms of *N*-glycans identified on the S-layer proteins from *S. tokodaii*. Multiple isomers of each glycoforms were also observed. The structure, branching and linkage of *N*-glycans were characterized by MS^n^ fragmentation by ESI-MS^n^ and linkage analysis by GC-MS.

**Figure S7:** *N*-linked glycosylation sites identified from SlaA of *S. tokodaii* by LC-MS/MS analysis (Tryptic digestion and semi-specific cleavage search using Byonic software).

(A) HCD MS^2^ spectra of glycopeptide ^1175^IYY<u>**N[SuphoQuinovose_1_Hex_4_HexNAc_2_] AT**</u>SGR^1183^ from SlaA.

(B) HCD MS^2^ spectra of glycopeptide ^1188^NVYGQVVL<u>**N[SuphoQuinovose**_**1**_**Hex**_**4**_**HexNAc**_**2**_**]AS**</u> GN^1200^ from SlaA.

(C) HCD MS^2^ spectra of glycopeptide ^1222^AVLP<u>**N[SuphoQuinovose_1_Hex_4_HexNAc_2_]NT**</u>LTTL TFNK^1236^ from SlaA.

(D) HCD MS^2^ spectra of glycopeptide ^1298^IIPA<u>**N[SuphoQuinovose_1_Hex_4_HexNAc_2_]IT**</u>PIR^1307^ from SlaA.

(E) HCD MS^2^ spectra of glycopeptide ^1362^EGV<u>**N[SuphoQuinovose_1_Hex_4_HexNAc_2_]AS**</u>VTSPV VYYSYQAV VAK^1383^ from SlaA.

(F) HCD MS^2^ spectra of glycopeptide ^1421^AVGPAISEYPVNLVFT<u>**N[SuphoQuinovose_1_Hex_4_ HexNAc_2_]VT**</u> VEK^1442^ from SlaA.

**Figure S8:** N-linked glycosylation sites identified from SlaB of *S. tokodaii* by LC-MS/MS analysis (Tryptic digestion and semi-specific cleavage search using Byonic software).

(A) HCD MS^2^ spectra of glycopeptide ^200^G<u>**N[SuphoQuinovose_1_Hex_4_HexNAc_2_]QT**</u>ISLTLK^209^ from SlaB.

(B) HCD MS^2^ spectra of glycopeptide ^347^EIETV<u>**N[SuphoQuinovose_1_Hex_4_HexNAc_2_]QT**</u>VYTL MNEIK^363^ from SlaB.

(C) HCD MS^2^ spectra of glycopeptide ^364^SL<u>**N[SuphoQuinovose_1_Hex_4_HexNAc_2_]AS**</u>ISQLSTTL SSTTTEITTLE NDIK^392^ from SlaB.

